# Phase-specific RNA accumulation and duplex thermodynamics in multiphase coacervate models for membraneless organelles

**DOI:** 10.1101/2021.05.15.444314

**Authors:** Saehyun Choi, Philip C. Bevilacqua, Christine D. Keating

## Abstract

Liquid-liquid phase separation has emerged as an important means of intracellular RNA compartmentalization. Some membraneless organelles host two or more compartments serving different putative biochemical roles; the mechanisms for, and functional consequences of, this subcompartmentalization are not yet well understood. Here, we show that adjacent phases of decapeptide-based multiphase model membraneless organelles differ markedly in their interactions with RNA. Additionally, their coexistence introduces new equilibria that alter RNA duplex stability and RNA sorting by hybridization state. These effects require neither biospecific RNA binding sites nor full-length proteins. As such, they are general and point to more primitive versions of mechanisms operating in extant biology that could aid understanding and enable design of functional artificial membraneless organelles.

---

Coacervates formed by liquid-liquid phase separation are important model systems for many membraneless organelles found in living cells.^1–3^ Since RNAs play critical roles in both the formation and biological functions of intracellular membraneless organelles,^4–6^ the effect of compartmentalization in coacervates on RNA distribution, structure and bioactivity is of particular importance. Accumulation of RNA into coacervates^7–11^ or membraneless organelles^4,5,12^ can localize and enhance its activity, for example facilitating ribozyme reactions. Nott and coworkers showed that nucleic acid partitioning in protein-based coacervate droplets depended on the nucleic acid length and hybridization levels.^13,14^ Additionally, DNA duplex dissociation was favored in the droplets as compared to the external dilute phase.^13^ In vivo experiments have demonstrated that RNA binding proteins (RBPs), constituting membraneless organelles, can make RNAs less-structured^15,16^ and less-entangled^17^, whereas the knockdown of abundant RBPs led to increased levels of double-stranded RNAs^18^. With these types of observations, we begin to glimpse mechanisms by which cells could take advantage of liquid-liquid phase separation in organizing RNA biochemistry.

Several membraneless organelles including nucleoli^12,19,20^, stress granules and P granules^21,22^ are themselves subdivided into two or more compartments, with different phases thought to perform distinct functions relating to, for example, RNA processing and gene expression.^6,12,23^ The phases have distinct protein compositions, and localize different types of RNAs.^12,23^ In nucleolar multiphase compartments, for example, nascent ribosomal RNAs (pre-rRNA) are located in the inner, FBL-rich phase where pre-rRNA processing and modification occur.^12,20^ On the other hand, rRNAs are located in the outer, NPM1-rich phase where they form complexes with ribonucleoproteins.^12,24^ FBL proteins have arginine-rich sequences that interact with pre-rRNAs and promote their accumulation in inner phase^20^, and NPM1 proteins have acidic and basic amino acid-rich sequences^25^, while both have RNA binding domains.^12,25^ More generally, individual phases of multiphase organelles likely take advantage of differences in RNA length- and structure-dependent interactions with their protein components to control RNA distributions and spatially organize biochemical functions.^12^

In an important step towards capturing this aspect of cell biology, multiphase coacervate model systems have recently been reported.^1–3,26–30^ Coexisting phases of these multiphase coacervates exhibited different local viscosities^26^ and partitioning behaviors.^1,3,29^ In coexisting coacervate systems, phases enriched in molecules better able to participate in base-pairing or cation-pi interactions accumulated simple single-stranded oligoRNA homopolymers (uridylic acid or adenylic acid 15-mers) more effectively^1^. However, it is not yet known how general these observations are across RNAs of other strandedness or sequence, nor how the ability of multiphase coacervates to control spatial distributions of encapsulated RNAs influences functional aspects such as their dissociation thermodynamics.

Herein, we demonstrate that the phases in multiphase complex coacervate droplets formed by combining three simple oligopeptides affect RNA chemistry via differences in RNA-peptide interactions. The adjacent phases within these droplets have remarkably different partitioning of oligonucleotides, with the inner phase preferentially accumulating single-stranded (ss) RNA, while the outer phase preferentially accumulates double-stranded (ds) RNA. Greater duplex dissociation (“helicase activity“) is also observed in the inner phase of the multiphase coacervate droplets. These differences between adjacent coacervate phases exceed those for single-phase coacervate droplets produced from pairwise peptide mixtures. This work shows that distinct phases within multiphase complex coacervate droplets not only recruit RNAs differently depending on their hybridization status, but also alter the RNA hybridization levels via a mechanism that takes advantage of the coupled partitioning and dissociation equilibria in the multiphase systems.

## Results and Discussion

### Multiphase droplet formation by oligopeptides

We chose the decapeptides of arginine (R10), lysine (K10), and aspartic acid (D10) as minimal models for key types of interactions leading to coacervation. These amino acids are common in repeating sequences of phase separating proteins in cells.^12,31^ Their charged sidechains provide ion pairing interactions that drive complex coacervation between the cationic R10 or K10 and the anionic D10. Both cationic peptides are expected to interact with RNAs via ion pairing with the backbone and cation-pi interactions with the nucleobases, with R10 expected to interact more strongly due to its more favorable cation-pi and pi-pi binding with the nucleobases.^32,33^ In comparison to longer peptides (e.g., 100mers), shorter peptides such as these have shown stronger partitioning of oligoRNA^7^ and could be more readily engineered to control properties of coacervate droplets for future application. Peptides were combined pairwise (R10/D10 and K10/D10) and all together (R10/K10/D10) to form coacervate droplets (Fig. 1 and Supplementary Table 1). Droplets formed in all three systems, but the R10/K10/D10 droplets were multiphase, with an inner coacervate phase entirely surrounded by an outer coacervate phase in each droplet (Fig. 1). This core-shell morphology indicates a greater interfacial tension between the inner phase and the dilute continuous phase than that between the outer coacervate phase and the continuous phase, suggesting higher overall peptide content (lower water content) for the inner phase.^34^ Direct measurements of peptide concentrations (next section) confirm this notion.

**Fig. 1.**
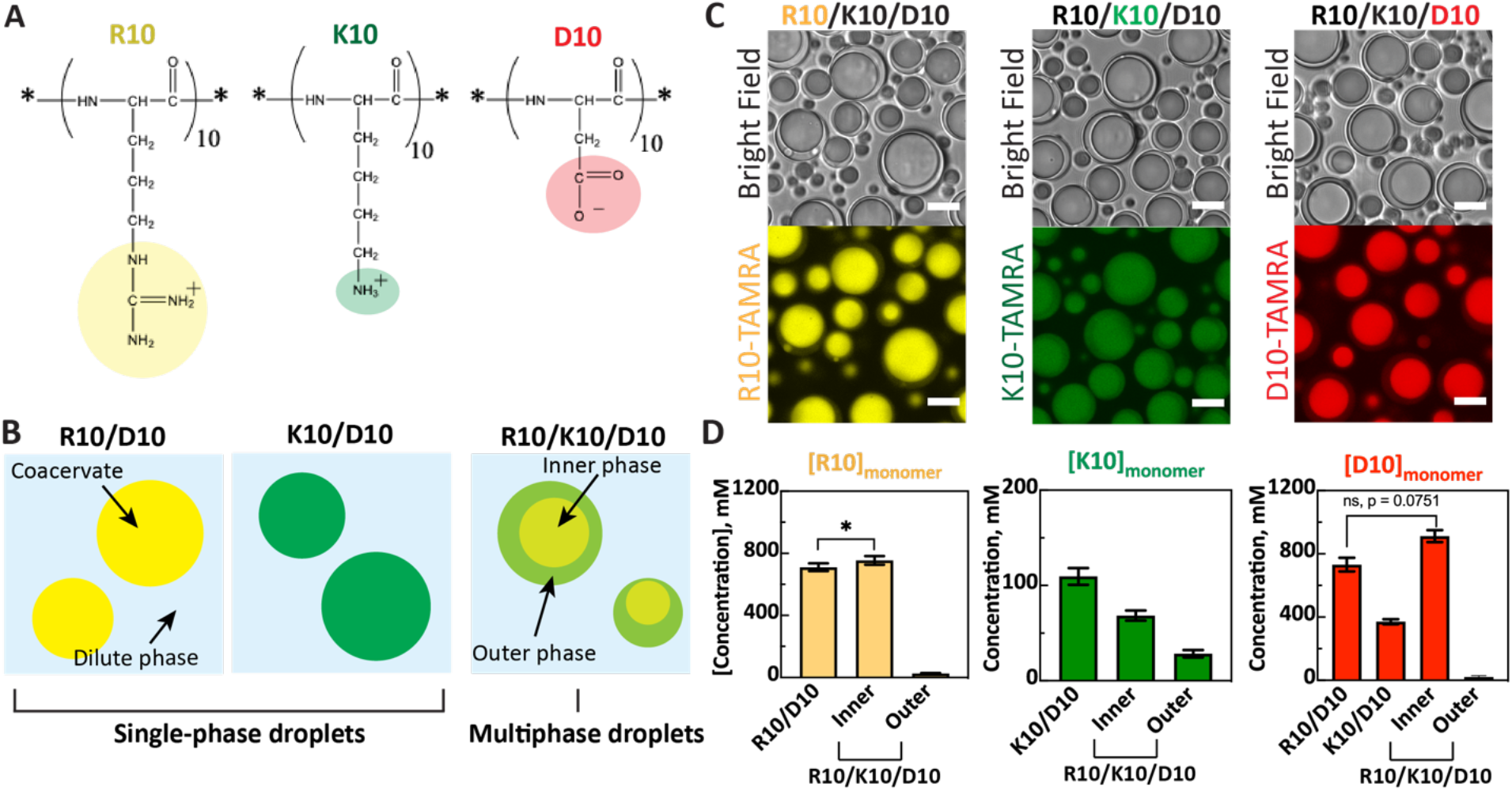
Formation of multi- and single-phase oligopeptide coacervates. **(A)** Molecular structure of the two cationic peptides, K10 and R10, and the anionic D10. **(B)** Illustration of coacervate droplets formation by mixing D10 with cationic peptides, together (left), separately to form single-phase droplets or (right), together to form multiphase droplets. The color of the cation from panel A is used to color each droplet. **(C)** Microscope images of multiphase R10/K10/D10 coacervate droplets, with each peptide fluorescently labeled to determine its distribution. Scale bars = 10 μm. **(D)** Quantification of local concentrations of R10, K10 and D10 sidechains in single-phase and multiphase coacervate droplets. “Inner” and “Outer” indicate the respective phases of the R10/K10/D10 coacervate droplets. All peptide coacervate samples were prepared in 15 mM KCl, 0.5 mM MgCl_2_ pH 8.3 ± 0.1, at 40 mM in charge-matched monomeric units. This corresponds to 40 mM positively-charged moieties (from 4 mM R10, or 4 mM K10, or 2 mM R10 plus 2 mM K10), mixed with 40 mM negatively-charged moieties from 4 mM D10. Error bars are standard deviation of ~45 samples from three independent trials. Two-sided t-tests with unequal variance were performed; all p-values are in Supplementary Table 3 and some p-values are included to aid comparisons. *: p-value < 0.05 and ns: statistically non-significant values.

### Peptide quantification in each coacervate phase

Concentrations of R10, K10 and D10 in each phase for R10/D10, K10/D10 and R10/K10/D10 coacervate droplets were determined in parallel experiments using TAMRA-labeled versions of each peptide (Fig. 1D and Supplementary Table 1, 2). Notably, the two phases of the multiphase coacervates do not correspond to a simple coexistence of phases formed by each cation/anion pair. We found that the inner coacervate phase of R10/K10/D10 has higher concentrations of *all three* oligopeptides than the outer coacervate phase (Fig. 1D). Local R10 and D10 concentrations in the inner coacervate phase are similar to those found in R10/D10 coacervates, while K10 is also present but at slightly lower concentration. The surrounding outer coacervate phase has roughly equal concentrations of all three peptides, all of them much more dilute than in the inner phase (Fig. 1D). Coalescence of multiphase R10/K10/D10 coacervates shows rapid fusion of the outer coacervate phases, followed by a slower fusion of the inner coacervate phases (Supplementary Fig. 1); this observation indicates lower viscosity for the outer coacervate phase consistent with its lower peptide content.^26^ Interestingly, while both R10/D10 coacervates, and the inner coacervate phase of R10/K10/D10 have cationic to anionic peptide ratios near 1, neither K10/D10 coacervates nor the outer coacervate phase of R10/K10/D10 are charge-balanced in terms of peptide sidechain occupancy (Supplementary Table 2 and Supplementary Discussion 1). Increasing salt concentration results in loss of the outer coacervate phase, consistent with differences in its peptide composition (Supplementary Fig. 2 and Supplementary Discussion 2).

### RNA partitioning in multiphase droplets

Interactions with peptides in the coacervate phases can be expected to drive accumulation of RNA and impact its structure, for example its ability to form base pairs and RNA duplexes. We therefore characterized RNA oligonucleotide accumulation in each coacervate phase for R10/D10, K10/D10, and R10/K10/D10 systems. Specifically, we evaluated single-stranded (ss) and double-stranded (ds) versions of 10-mer and 20-mer sequences designed to avoid self-complementarity and to have the same GC content when double stranded (Supplementary Table 4). These dsRNAs were confirmed to form duplexes in 15 mM KCl, 0.5 mM MgCl_2_ and pH 8.3 ± 0.1 by the presence of cooperative melting transitions in UV-Vis melting experiments that had expected increases in stability with duplex concentration and salt concentration (Supplementary Fig. 5). For all RNAs tested and all coacervate phases, RNA partitioned strongly into the droplets, reaching concentrations from 20-to 700-fold higher than the level expected if the RNA distributed equally across coacervate and dilute phases (0.1 μM) (Fig. 2). However, there were also considerable differences between the phases and the different RNAs.

**Fig. 2.**
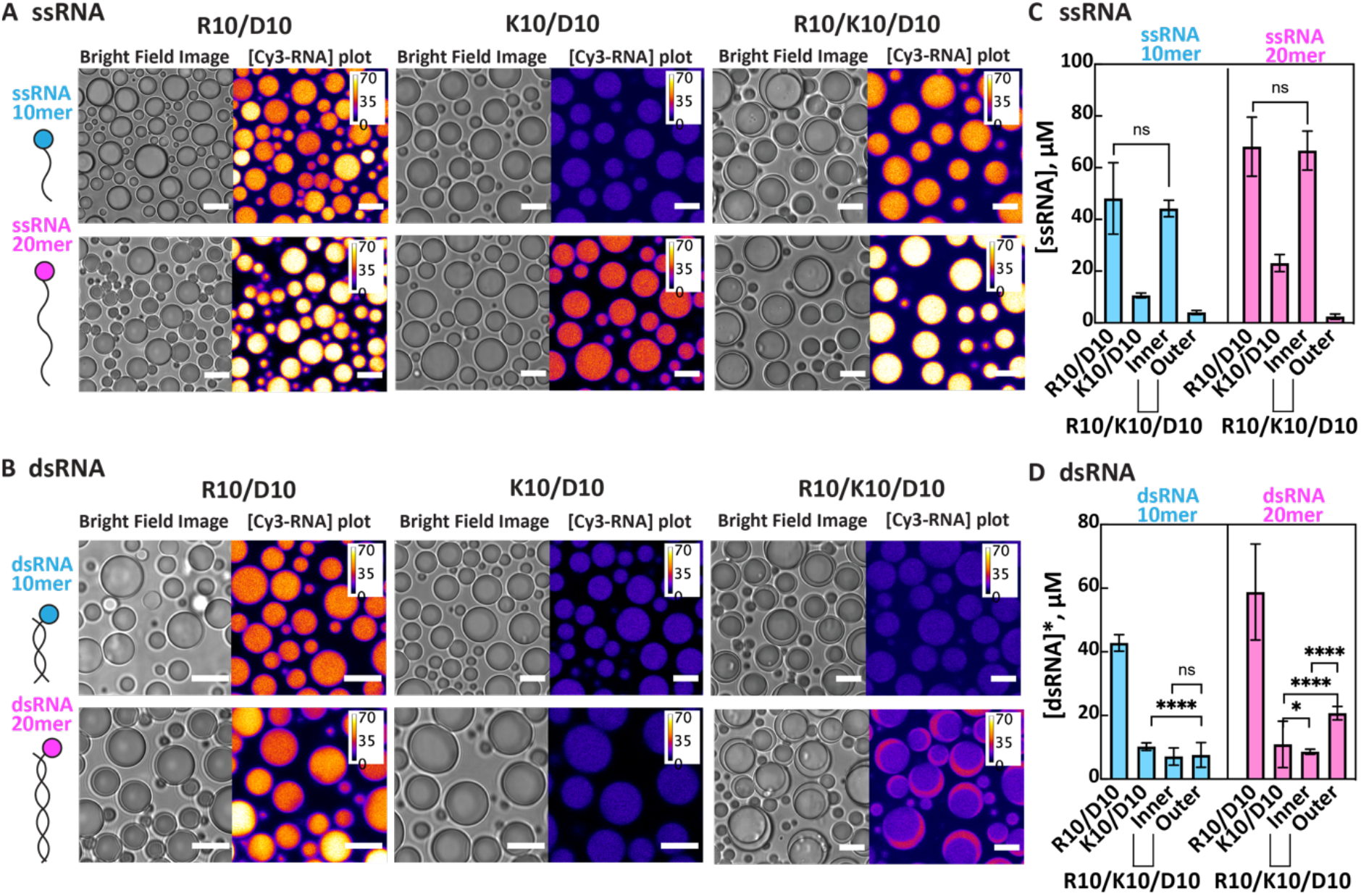
Partitioning of RNA in single- and multiphase peptide coacervates. Pairs of bright field images (left) and [Cy3-RNA] plots (right) of **(A)** ssRNA and **(B)** dsRNA in R10/D10, K10/D10 and R10/K10/D10 droplets. Scale bars = 10 μm. Calibration bars on [Cy3-RNA] plot in μM converted from the intensity plot using the calibration curves of various concentration of Cy3-RNA solution. **(C, D)** Bar graphs of average concentration of Cy3-RNA in each phase for **(C)** ssRNA ([ssRNA])and **(D)** dsRNA ([dsRNA]*). [dsRNA]* in dsRNA partitioning data assumes that prehybridized RNA remained in duplex form; however, due to possible re-equilibration within coacervate phases, a portion of the signal could also arise from ssRNA. Either ssRNA or dsRNA were added at a final concentration of 0.1 μM to each sample. The dsRNAs were prepared by mixing 5’-Cy3 labeled ssRNA (0.1 μM) with its ssRNA complement (0.1 μM). Fluorescence images of Cy3-RNAs and local RNA concentration values can be found in Supplementary Fig. 4 and Supplementary Table 5. Error bars show standard deviation of measurements of ~ 45 samples from three independent trials. Two-sided t-test with unequal variance was performed for all pairs of data sets and all p-values are available in Supplementary Table 6, 7 and 8; some p-values are included in panels C and D to aid comparisons. *: p-value < 0.05, ****: p-value < 0.0001, ns: not significantly different values.

Single-stranded RNA accumulated preferentially in the inner coacervate phase of the R10/K10/D10 droplets, with 10- and 20-mer ssRNAs reaching local concentrations ~10-fold and ~30-fold higher than in the outer coacervate phase of these droplets and up to ~660-fold higher than the 0.1 μM overall concentration added (Supplementary Table 5). The R10/D10 single-phase coacervates had similar partitioning of ssRNAs as the inner coacervates, while K10/D10 single-phase coacervates had RNA accumulation intermediate between that of the inner and outer coacervates of R10/K10/D10. These RNA partitioning results can be interpreted in terms of the local availability of cationic sidechains in each phase: highest in R10/D10 and the inner phase of R10/K10/D10, lower in K10/D10, and lowest in the outer phase of R10/K10/D10 (Fig. 1D, Supplementary Table 2). K and R sidechains can interact with the phosphate backbone of ssRNAs via ion pairing and with the nucleobases via cation-pi, pi-pi, and hydrogen bonding interactions. Although anionic sidechains of D10 are also most concentrated in the inner coacervate phase of R10/K10/D10 and will compete with RNA for binding to R and K sidechains, the ssRNA has stronger interactions and is expected to displace the carboxylate moieties of D10.^9,35^

In contrast to ssRNA, dsRNA accumulated preferentially (~2.4x) in the outer coacervate phase over the inner coacervate phase of R10/K10/D10 (for the 20mer) or partitioned evenly between the inner and outer phases (for the 10mer) (Fig. 2B, D). The relatively poor accumulation of double-stranded RNAs into the inner coacervate phase of R10/K10/D10 is not expected based on the observed strong accumulation of dsRNA into R10/D10 single-phase coacervates, which have similar peptide composition to the inner coacervate phase of the multiphase droplets. Indeed, since the primary mechanism for RNA accumulation in any of the coacervate phases relies on interactions with the R10 and K10 peptides, it is perhaps surprising that dsRNA does not follow the same trend as ssRNAs with greater partitioning into the phases most enriched in cationic moieties. Reduced accessibility of its nucleobases in dsRNA as compared with ssRNA may inhibit cation-pi and pi-pi interactions with the peptides. Double-stranded oligonucleotides have been reported to partition less effectively in coacervates of the intrinsically disordered protein, Ddx4, which was interpreted as a consequence of their greater persistence length as compared to single-stranded oligonucleotides.^7,13^ Persistence length could be a factor here as well. The outer R10/K10/D10 coacervate phase may have a more open, dynamic mesh-like structure at the nanoscale as compared with the inner phase, consistent with its reduced peptide density and local viscosity.^13,26^ Additionally, the outer coacervate phase has an ~2.6x excess of cationic over anionic sidechains, which could enable dsRNA, having higher charge density than ssRNA^36^, to accumulate there despite competition with the D10 for interactions with the R10 and K10 (Supplementary Table 2).

### Phases of multiphase droplets impact duplex thermodynamics differently

Since RNAs distribute across the coacervate phases of R10/K10/D10 based on strandedness as demonstrated in the previous section, it is possible that RNA dissociation equilibria could be shifted. Dissociation of pre-hybridized dsRNA would cost ion pairing multivalency, but gain other interaction modes with the coacervate components, especially the guanidinium groups of R10, and decrease RNA's persistence length. We therefore sought to determine the level of dsRNA hybridization in each phase by Förster resonance energy transfer (FRET). In these experiments, a 3’-Cy3-labeled RNA is pre-hybridized with its antisense sequence having a 5’-Cy5-label at the same end of the duplex; these sequences were pre-hybridized in the same way as for the partitioning experiments described above.

We first consider the single-coacervate systems K10/D10 and R10/D10. The FRET results are summarized in Fig. 3 and Supplementary Fig. 6. Controls in buffer show a range of FRET efficiencies from near zero for single-stranded RNA to ~0.7 and ~0.8 for 10-mer and 20-mer duplexes, respectively (Fig. 3B). In K10/D10 coacervates, FRET efficiency for RNA 10-mer hybridization was similar to its value in buffer, while 20mer RNA showed ~20% lower FRET efficiency, indicating some dehybridization. The R10/D10 coacervates destabilized both 10-mer and 20-mer duplexes, with FRET values of ~0.5 and ~0.7, corresponding to roughly 20% decreases from each value in buffer. Such destabilizations have been reported previously for peptide- or protein-based condensates with respect to their surrounding dilute continuous phases, and are interesting as a physical form of “helicase activity”.^7,13^ For example, Nott et al reported that FRET efficiency decreased by 33% (to ~0.25) as compared to values in the dilute phase for 12mer or 24mer DNA duplex in droplets composed of the intrinsically-disordered protein, Ddx4.^13^ We previously found a length-dependence to this effect in K_n_/D_n_ peptides, with K30/D30 and K100/D100 coacervates destabilizing an RNA 10-mer, while K10/D10 had much less effect.^7^ Although the role of peptide or protein sequence in a coacervate's ability to melt duplexes has not yet been explored, it is reasonable to expect stronger destabilization for coacervates whose component molecules interact more strongly with RNA (or DNA), and particularly with its single-stranded forms. The greater destabilization we observe for R10/D10 as compared to K10/D10 is consistent with stronger interactions between the nucleobases and the guanidinium groups of R as compared with the amines of K,^30,32^ and with the observed stronger partitioning of ssRNA into the R10/D10 coacervates as compared with the K10/D10 coacervates (Fig. 2C).

**Fig. 3.**
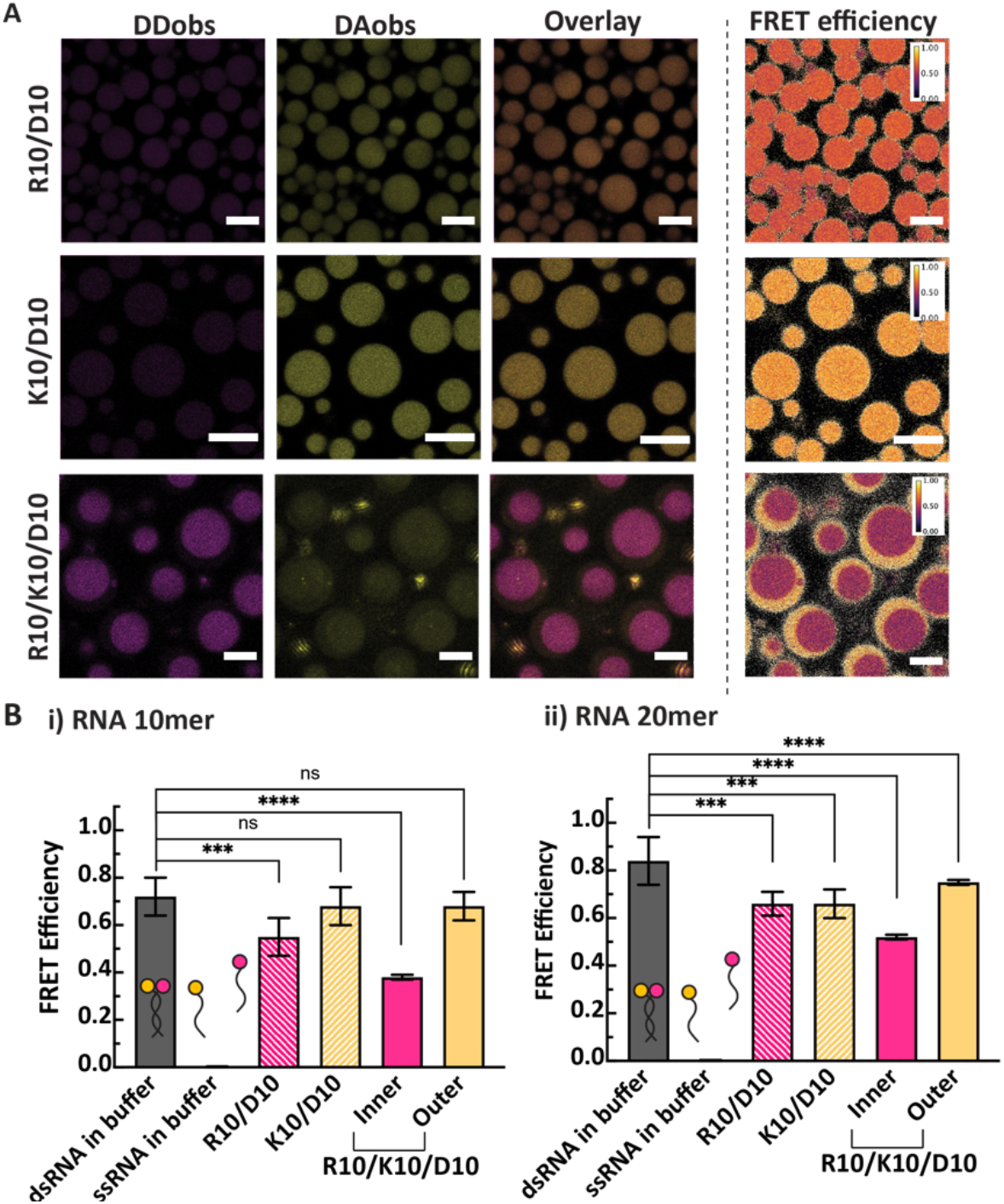
Comparison of RNA duplex stability in single- and multiphase peptide coacervates. **(A)** Fluorescence images and estimated FRET efficiency plot of dsRNA 10mer in R10/D10, K10/D10 or R10/K10/D10 coacervates. Scale bars = 10 μm. **(B)** FRET efficiency of i) RNA 10mer and ii) RNA 20mer in the same series of coacervate droplets. FRET of dsRNAs and ssRNA in buffer are controls that contain RNAs without polypeptides in solution. Cy3 is depicted in gold and Cy5 in red balls. All FRET efficiency values are shown in Supplementary Table 10. Error bars correspond to standard deviation of measurements of 9 samples from three independent trials. Two-sided t-tests with unequal variance were performed for FRET efficiency in coacervate phase to dsRNA in buffer for comparison. ****: p-value < 0.0001, ***: p-value < 0.001, and ns: not significantly different values. P-values of all tested pairs can be found in Supplementary Table 11. All peptide coacervate samples were prepared in 15 mM KCl, 0.5 mM MgCl2 pH 8.3 ± 0.1, at 40 mM in charge-matched monomeric units (Supplementary Table 1).

Turning now to the multiphase R10/K10/D10 coacervate system, we find relatively high FRET efficiencies for both lengths of RNA in the outer coacervate phase (~0.7 and ~0.8 for RNA 10mer and 20mer, respectively), similar to the FRET values in buffer. This is consistent with the lower peptide concentration of the outer phase. In contrast, we find that the inner phase of R10/K10/D10 coacervate droplets significantly destabilizes RNA duplexes, up to 46%. This phase has lower FRET efficiency than that of either type of single-phase coacervates for both RNAs, especially 10mer duplex consistent with ease of disrupting its intrinsically weaker structure. The two coacervate phases of R10/K10/D10 maintain substantially different environments for RNA duplex destabilization in adjacent compartments. FRET efficiency decreases by 44% (for 20-mer RNA) and 27% (for 10mer) from the outer to the inner coacervate phases, which is as large of a difference as is generally seen between a coacervate phase and its dilute supernatant phase^7,13^. It is notable that we can achieve this much difference in FRET efficiency between phases *within* individual droplets, particularly while single-phase droplets formed from the same peptides do not exhibit as large of destabilization. Special features of the multiphase coacervates include (1) their different phase compositions, with all three peptides distributed across both phases possibly destabilizing duplexes in a cooperative fashion, and (2) RNAs partitioning not only between the dilute phase and the coacervates, but also between the coexisting coacervate phases. Peptide composition alone does not offer an easy explanation of our observations. For example, local R10 concentrations in the inner coacervate phase of R10/K10/D10 are quite similar to R10/D10 (Figure 1D), but the inner phase of multiphase coacervates was notably more destabilizing for RNA duplexes, especially 10mer (Figure 3B). We next consider the thermodynamic balance between partitioning and dissociation in multiphase coacervates.

### Thermodynamic analysis for coupling RNA dissociation with partitioning between phases in multiphase droplets

In the absence of coacervation, cationic peptides are known to stabilize nucleic acid duplex formation.^37–40^ We verified this with our RNAs in low salt concentration in presence of R10, K10, or D10 by UV-Vis spectrometry-detected melting experiments, and confirmed that the RNA duplex was thermally stabilized in presence of R10 or K10 (Supplementary Fig. 5C). Duplex stabilization by cationic peptides seems to contradict our observation of lower FRET efficiency in the inner coacervate phase of R10/K10/D10 as compared with the outer coacervate phase, since the inner phase has a much higher concentration of cationic peptides (see Fig. 1D). We thus sought to account for both RNA partitioning and RNA dissociation equilibria to understand our multiphase droplet results.

We began by using the FRET data to estimate local concentrations of ssRNA and dsRNA in each phase for the pre-hybridized dsRNA partitioning experiments. After confirming dsRNA in buffer conditions at room temperature are hybridized from UV-Vis melting experiments, hybridization fractions of dsRNAs were estimated from normalized FRET efficiency using dsRNA and ssRNA buffer values. We estimated the concentration of dsRNA and ssRNA as follows using the fraction of hybridization: [*dsRNA*] = *F*_*hyb*_ × [*dsRNA*]* and [*ssRNA*] = 2 × (1 − *F*_*hyb*_) × [*dsRNA*]*, where F_hyb_ is the estimated fraction of RNA hybridization using FRET efficiency normalized to that of dsRNA in buffer, and [dsRNA]* is the concentration of Cy3-RNA in prehybridized dsRNA partitioning experiments in Fig. 2. Accordingly, we can define the equilibrium constants (K) for RNA partitioning from outer to inner coacervate phases for ssRNA and dsRNA as 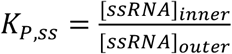 and 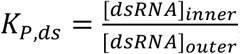, and K of RNA duplex dissociation in inner and outer phase as 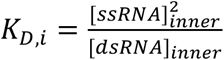 and 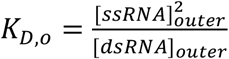 Upon correcting for partial melting of the dsRNAs, we see greater sorting of the duplexes to the outer than the inner coacervate phase of R10/K10/D10 with 2-fold and 3-fold greater concentration in the outer vs inner coacervate phase (K_P,ds_ ~ 0.5 and ~0.3) for the 10-mer and 20-mer, respectively than was apparent in Fig. 2 (Fig. 4, Supplementary Table 13, Supplementary Discussion 4).

**Fig. 4.**
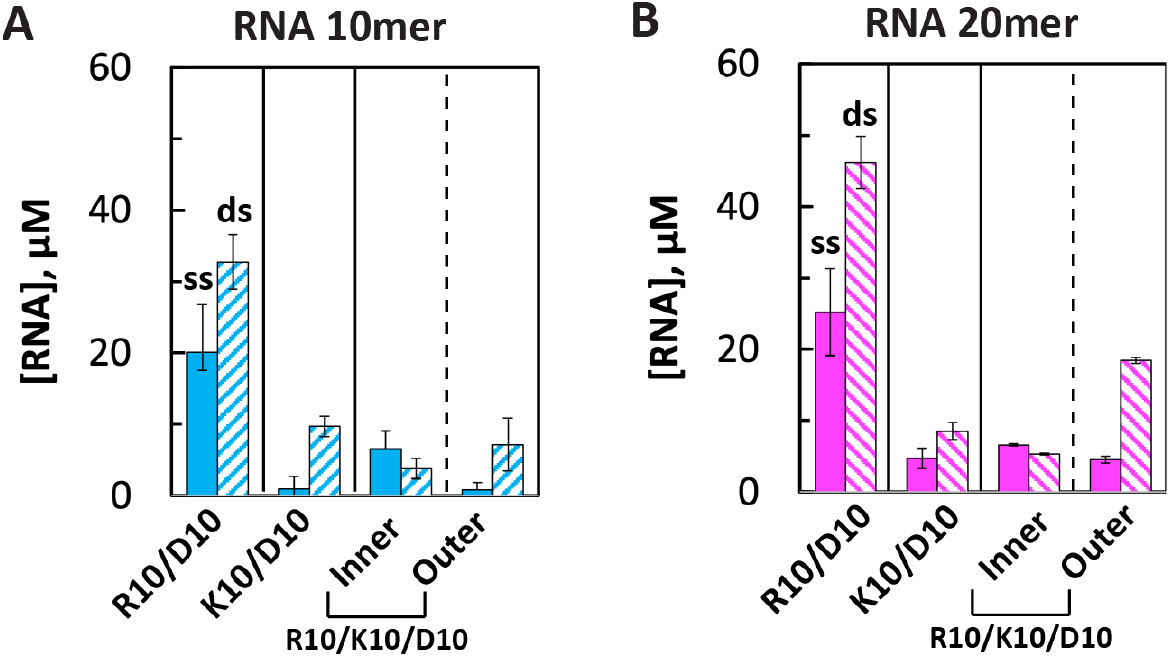
Estimated local concentrations of single- and double-stranded RNAs in each coacervate phase after equilibration of added dsRNA. Estimated total concentration of dsRNA (**ds,** [dsRNA]) and ssRNA (**ss**, [ssRNA]) from pre-hybridized dsRNA partitioning experiments in coacervate droplets in Figure 3 using the hybridization fraction estimated from FRET for **(A)** RNA 10mer and **(B)** RNA 20mer. Note that each dsRNA generates two ssRNAs upon dissociation. Error bars show propagated errors from standard deviation of partitioning experiments of 45 samples and FRET experiments of 9 samples of three independent trials.

Fig. 5A summarizes the coupled thermodynamic equilibria for RNA partitioning and dissociation in multiphase coacervates. We calculated Δ*G* for partitioning (Δ*G*_P_) of the ss and ds 10- and 20mer RNAs between two coacervate phases of R10/K10/D10 droplets (Δ*G*_P,ss_ and Δ*G*_P,ds_, respectively) based on Δ*G* = −*RTlnK*_P_. The Δ*G* for dissociation of RNA duplex in inner and outer phases, Δ*G*_D,i_ and Δ*G*_D,0_, were calculated based on Δ*G* = −*RTlnK*_D_ (see Methods for details). The dilute continuous phase between the droplets (not shown in Fig. 5A) has a much lower RNA concentration than either coacervate phase and is neglected in this analysis for simplicity. Estimated free energy values for partitioning based on the distribution of RNAs between the inner and outer coacervate phases are summarized for each length and strandedness in Fig. 5B. Single-stranded RNAs accumulated in the inner coacervate phase (Δ*G*_P,ss_ < 0), while double-stranded RNAs preferentially accumulate in the outer coacervate phase (Δ*G*_P,ds_ > 0). This effect is stronger for the shorter ssRNA 10-mer as compared to the ssRNA 20-mer (Fig. 5C). The dissociation free energy is always positive (Δ*G*_D,i_ > 0 *and* Δ*G*_D,0_ > 0), reflecting the favorability of duplex formation for the complementary RNAs (Fig. 5D). We also see differences in dissociation free energy between the phases, with dsRNA more favored in the outer coacervate phase of the multiphase droplets (Fig. 5E).

**Fig. 5.**
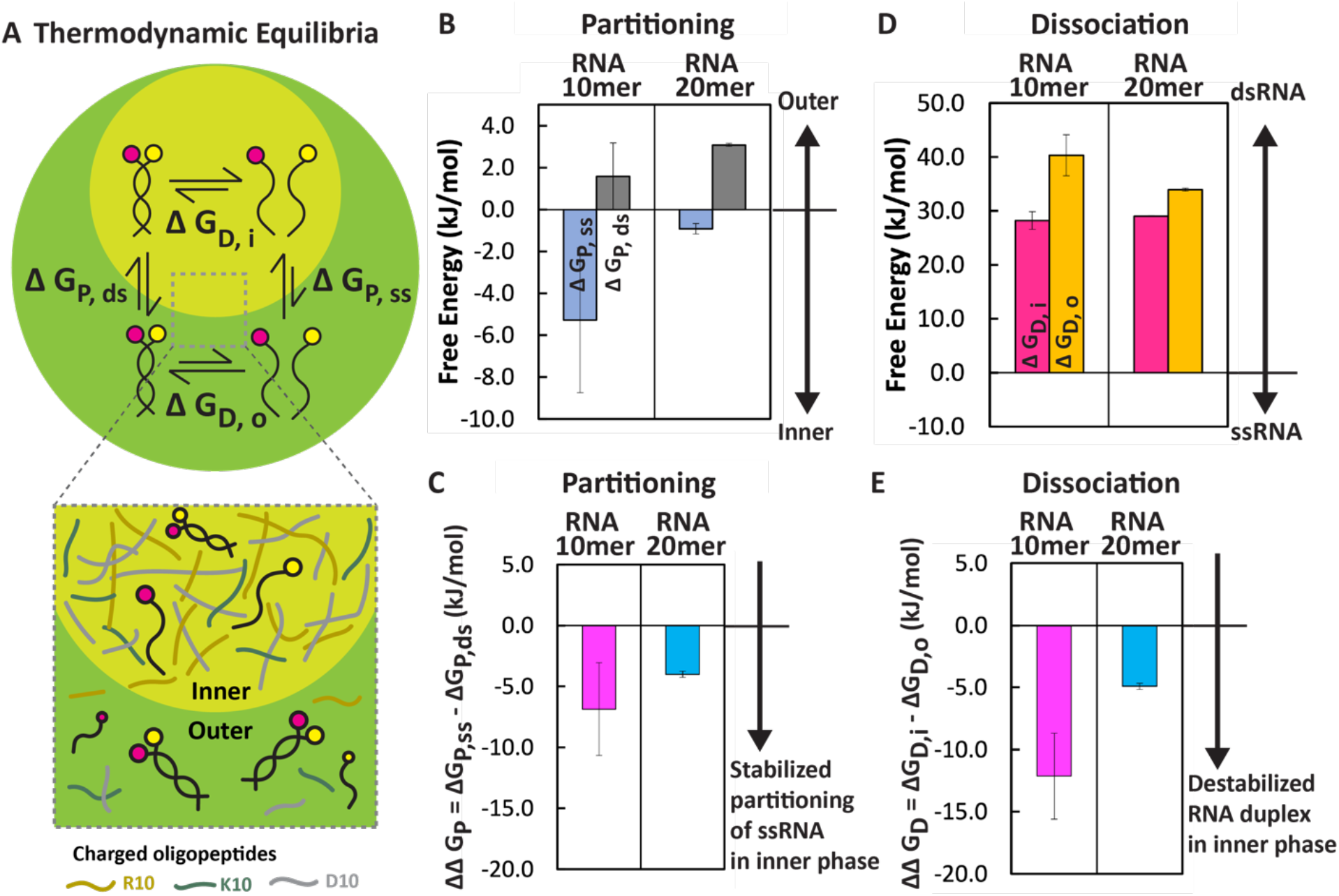
Coupling of partitioning and dissociation equilibria in multiphase coacervate droplets dictates distribution of single- and double-stranded RNAs. **(A)** Scheme of thermodynamic equilibrium of partitioning and dissociation of dsRNAs in multiphase droplets. Although some RNA also exists in the dilute phase that surrounds the droplets, is quite dilute and its contribution is neglected in this analysis. **(B)** Gibbs free energy change in partitioning of dsRNA (Δ*G*_P,ds_) and ssRNA (Δ*G*_P,ss_) from outer to inner coacervate phase. **(C)** Stability of partitioning of ssRNA into inner phase (ΔΔ*G*_P_) calculated from change between Δ*G*_P,ss_and Δ*G*_P,ds_in (B). **(D)** Gibbs free energy change in dissociation of dsRNA into ssRNA in inner (Δ*G*_D,i_) and outer droplets (Δ*G*_D,0_). **(E)** Stability of RNA duplex in inner phase (ΔΔ*G*_D_) calculated from the difference of Δ*G*_D,i_ and Δ*G*_D,0_ in (C). Free energy of partitioning and dissociation is calculated from equilibrium constants using estimated concentrations of ssRNA and dsRNA in each phase from Fig. 4 (see Methods for details). Error bars show propagated errors from standard deviation of partitioning experiments of 45 samples and FRET experiments of 9 samples of three independent trials; all values are in Supplementary Table 12.

Interestingly, the dissociation free energies for RNA duplex in the inner and outer R10/K10/D10 coacervate phases were similar to values for single-phase droplets of R10/D10 and K10/D10, respectively (Supplementary Fig. 7C), although both single-coacervate systems showed only slight dsRNA dissociation in FRET experiments (Fig. 3). Additional thermodynamic equilibria present only for the multiphase coacervate samples can explain this nonintuitive result. Considering the differences in Δ*G* for each equilibrium, ssRNA accumulation into the inner coacervate phase is stabilized as shown by the difference of the partitioning free energy of ssRNA and dsRNA (Fig. 5C, ΔΔ*G_P_* < 0), while RNA duplex is destabilized in inner phase according to the difference in the dissociation free energy between inner phase and outer phase (Fig. 5E, ΔΔ*G*_D_ < 0)). Although the magnitudes for the dissociation free energies are substantially larger than the partitioning free energies (Fig. 5B, C), their differences are of a similar magnitude (Fig. 5C, E, compare between same colors). These similarities, which represent subtractions of the opposite edges of the thermodynamic box in Fig. 5A, also support the validity of the calculations since all four values were determined independently. The favorable partitioning of ssRNA into the inner phase can explain the observed higher levels dsRNA dissociation in inner phase. We conclude that the coupling of partitioning and dissociation equilibria between adjacent coacervate phases is critical to the strong RNA helicase-like activity of the inner coacervate phase seen in Fig. 5E.

## Conclusion

The work presented here demonstrates that multiphase coacervate systems are more than the sum of their component phases - their coexistence introduces new partitioning equilibria important in controlling the distribution of RNAs in different hybridization states. The differences we observed between multiphase R10/K10/D10 coacervates and related single-coacervate systems (R10/D10 and K10/D10) go beyond the partial sharing of polyelectrolytes across phases that has been reported previously^41^. First, the outer coacervate phase of the R10/K10/D10 system is substantially more dilute in total peptides than either of the single-coacervate systems, and particularly lacks anionic moieties. Second, coupling of equilibria for RNA partitioning and RNA dissociation in each phase of multiphase coacervates determines the spatial distribution of duplex destabilization. Differences in RNA partitioning and dissociation thermodynamics between the two coexisting coacervate phases of R10/K10/D10 are consequently larger than those between R10/D10 and K10/D10 coacervates. Our findings in these simple oligopeptide systems point to potential underlying mechanisms by which nucleic acids can be sorted based on their hybridization state, and their duplexes can be stabilized or destabilized locally within a multiphase membraneless compartment even in the absence of specific binding motifs or full-length proteins.

## Methods

### Quantification of local R10, K10 and D10 concentration

Fluorescent-labeled peptide stock solutions were mixed with coacervate samples to be the final concentration as stated in Supplementary Table 1. TAMRA-labeled peptides were mixed with non-fluorescently labeled peptides to be 0.5 % or 0.25 % by concentration (Supplementary Table 1). We used low percentage concentration of fluorescently-labeled peptides, so that we can minimize artifacts in coacervate phases^1,41^. Intensity of labeled polypeptides were averaged from 45 droplets total from three independent trials after the background correction. Calibration curves of fluorescent labeled R10, K10 and D10 are acquired with the same confocal microscope setting as a function of fluorescent labeled R10, K10 and D10 concentrations over fluorescence intensity. Stock solutions of fluorescent labeled R10, K10 and D10 in water were used for the calibration by diluting it with water. Using the calibration curves, we calculated the concentration of fluorescent labeled R10, K10 and D10, and then, used the dilution factor to estimate the monomer concentration of R10, K10 and D10 (Supplementary Table 1).

### Quantification of RNA concentration

Final total concentration of RNA is 0.1 μM for all samples. Pre-hybridized dsRNAs were prepared in 150 mM KCl after melting in 90 °C for 2 mins with following equilibration at room temperature for 1 hour to allow hybridization before they were mixed into coacervate samples. Calibration curves as a function of concentration of fluorescent-labeled RNAs over fluorescent intensity were individually achieved with the same setting of confocal microscopes using various concentration of 5’-Cy3-ssRNA 10mer, 5’-Cy3-ssRNA 20mer, pre-hybridized dsRNA 10mer and pre-hybridized dsRNA 20mer from 40 μM to 1 μM.

### FRET efficiency calculation and the fraction of hybridization to estimate concentration of ssRNA and dsRNA in pre-hybridized dsRNA partitioning

We used the methods published in literature by Nott et al.^13^ and Cakmak et al.^7^ For Föster resonance energy transfer (FRET), 3’-Cy3-ssRNA and its complementary sequence of 5’-Cy5-ssRNA were used. Cy3 is used as donor with 543 nm laser excitation and its emission was collected between 555-625 nm. Cy5 is the acceptor with 633 nm laser excitation and its emission was collected between 650 nm-750 nm. Three channels of fluorescence images were as following: DD_obs_ (Donor emission after donor excitation), DA_obs_ (Acceptor emission after donor excitation), AA_obs_ (Acceptor emission after acceptor excitation). The emission and absorbance wavelengths of donor and acceptor dyes can overlap, so they should be corrected. The samples containing donor-only or acceptor-only dyes as stated in Table S4 were imaged using same parameters of confocal microscope to calculate two correction values: *α* and *β* respectively; *α* = *DA*_*donor*_/*DD_donor_*, *β* = *DA_acceptor_*/*AA_acceptor_*. The corrected FRET efficiency (ECT) is calculated using following equation

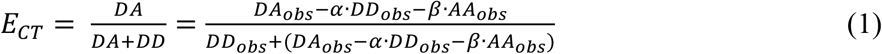

To calculated FRET efficiency, mean intensities are used from three samples prepared, and each sample has three images containing three droplets. FRET plots were processed from fluorescence intensity images using Fiji^42^ using averaged *α* and *β*. As controls, dsRNA in buffer and ssRNA in buffer were used; both controls contain 5 μM of each RNA strands as stated in Supplementary Table 4 with final concentration of 15 mM KCl, 0.5 mM MgCl2 and 10 mM Tris (pH 8.1). dsRNAs in this solution condition are shown to be hybridized from UV-Vis melting experiment, and thus, maximum FRET efficiency is expected (Supplementary Fig. 4). ssRNA buffer is using same sequence of RNAs each with Cy3 and Cy5 dye, so no hybridization will occur, which is shown as FRET efficiency close to zero. Because the ECT of dsRNA buffer and ssRNA buffer will be maximum and minimum respectively, we can estimate fraction of hybridization (Fhyb) by normalizing FRET efficiency of systems with FRET efficiency values of dsRNA buffer and ssRNA buffer as 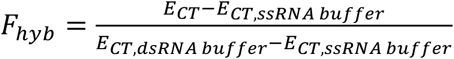, which is based on assumption made in analysis methods for FRET pairs for nucleic acid hybridization sensors^43,44^. When pre-hybridized dsRNAs are accumulated in coacervate droplets, it is expected that Cy3-RNA fluorescence intensity ([dsRNA]*in Fig. 2) is coming from the mixture of Cy3-RNA hybridized (dsRNA form, [dsRNA]) and dissociated (ssRNA form, [ssRNA]). Therefore, we estimated the concentration of dsRNA and ssRNA as following using the fraction of hybridization: [*dsRNA*] = *F*_*hyb*_ × [*dsRNA*]* and [*ssRNA*] = 2 × (1 − *F*_*hyb*_) × [*dsRNA*]*.

### Thermodynamic calculation of RNA partitioning and hybridization in multiphase droplets

We defined thermodynamic parameters using the thermodynamic equilibria noted in Fig. 4A and

Supplementary Fig. 7A. Equilibrium constants for each part is defined as 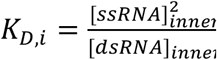 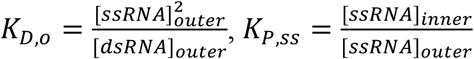and 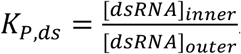 Gibbs free energy is calculated using Δ*G* = −*RT* ln (*K*), where R is gas constant and T is room temperature in Kelvin, for example, Δ*G*_*P,ss*_ = −*RTlnK_P,ss_*, Δ*G_P,ds_* = −*RTlnK*_P,ds_, Δ*G_D,i_* = −*RTlnK_D,i_* and Δ*G_D,0_* = −*RTlnK_D,0_*.

## Acknowledgments

This work was supported by the NASA Exobiology program grant 80NSSC17K0034. S.C. was supported by Future Investigators in NASA Earth and Space Science and Technology (FINESST) under Grant 80NSSC19K1531.

## Additional Information

Supplementary Information is available in the online version of the paper.

## Author Contributions

S. C. performed the experiments. All authors conceived and designed the experiments and analyzed the data. S. C. and C. D. K wrote the manuscript with input from P. C. B.

## Notes

### Competing Interest Statement

The authors have declared no competing interest.

