## Supplementary Information for "Phase-specific RNA accumulation and duplex thermodynamics in multiphase coacervate models for membraneless organelles"

**Materials and Supplementary Methods.**

**Chemicals.** Poly(L-Arginine hydrochloride) (degree of polymerization n = 10) (R10), Poly(L-Lysine hydrochloride) (degree of polymerization n = 10) (K10) and poly(L-aspartic acid sodium salt) (degree of polymerization n = 10) (D10) were purchased from Alamanda Polymers and used without further purification and they are 95% purity or above. N-terminus TAMRA labeled versions of R10, K10 and D10 were purchased from Biomatik. All oligoRNAs are custom-synthesized and purchased from Sigma Aldrich.

**Preparation of coacervate samples.** Peptide stock solutions were prepared as 10 mM final concentration in HPLC grade water and pH adjusted to pH 8.1-8.3 using 1 M NaOH or HCl. Aliquots of peptide stock solutions were sealed with inert gas in and kept under -22 °C for storage. Before use, peptide stock solutions were thawed and vortexed thoroughly. Coacervate samples are always freshly prepared before characterizations. We prepared samples with the following order of addition: HPLC water, salt, D10 stock solution, K10 stock solution, R10 stock solution, and fluorescent-labeled RNA or fluorescent-labeled peptide. Coacervate samples were equilibrated at room temperature for 15 mins before analysis. Polypeptides were mixed in 15 mM KCl and 0.5 mM $MgCl_2$ solution for all analysis, except salt dependent multiphase coacervate formation analysis in Fig. S1. Final concentration of peptides in coacervate samples is listed in Table S1.

**Instrumentation and image analysis.** Confocal images were taken using a Leica TCS SP5 laser scanning confocal inverted microscope (LSCM) with Leica LAS AF software and an HCX PL APO CS 63.0×/1.40 oil UV objective. Fluorescence intensity values were acquired from raw fluorescence images using Fiji[1] after background correction.

**Supplementary Discussion 1**

We quantified peptide concentrations using TAMRA N-labeled versions of each peptide in parallel experiments. Calibration curves for each TAMRA N-labeled peptide were individually obtained. The peptides used here are relatively low molecular weight, and some impact of labeling on peptide partitioning cannot be entirely ruled out; however, by using the same label for all peptides, this effect should be similar across the three peptides and still allow us to compare their relative distributions. We note that the attached TAMRA label is zwitterionic, so the net charges of TAMRA N-labeled polypeptides are unchanged.

It is interesting to compare the charge ratio of cationic to anionic moieties in each coacervate phase for the three systems. This ratio was close to 1:1 (balanced cationic and anionic moieties coming from the sidechains of cationic vs anionic peptides) for R10/D10 coacervates and for the inner coacervate phase of R10/K10/D10 (Supplementary Table 2). However, we observed a roughly threefold excess of anionic moieties in K10/D10 coacervates, and a more than twofold excess cationic moieties in the outer coacervate phase of R10/K10/D10. Although ratio of cationic groups and anionic groups in coacervate phases are generally expected to be close to 1 in polyelectrolyte-based coacervates[2], the imbalance of stoichiometry of polycations and polyanions in coacervate phase can be suggested by charged by zeta potential measurements of coacervate droplets[3] or highest optical turbidity of coacervate samples at non-stoichiometric ratio of polycation and polyanion[4]. The pH for our experiments is 8.3 +/- 0.1, which is close to the pKa values for the peptide N-terminal amines (pKa ~ 8) and the side chain of Lys (pKa ~ 10)[5], thus by assuming all peptide sidechains are charged, we may be overestimating the cationic moieties[4]. It should be noted that an imbalance of stoichiometry of cationic and anionic groups from the peptides can be compensated by free ions in each phase to maintain charge neutrality in phase[6], which we did not measure in this work. Of particular note in this context is $Mg^{2+}$, which was present at 0.5 mM and which can bind to the carboxylates of the oligoaspartates.

**Supplementary Discussion 2**

Since coacervates formed using polyR and polyK as the cationic species are known to have different salt stabilities[7,8], we tested the effect of increasing KCl concentration on multiphase droplet morphology. The R10/K10/D10 multiphase coacervates became single-phase droplets as salt concentration increases (Supplementary Fig. 2). It appears from the images of Supplementary Fig. 2 that the inner coacervate phase, which is particularly enriched in TAMRA-R10, persists while the outer coacervate phase dissolves. Local peptide concentrations in all phases also decreases with increasing KCl concentration, noted by the less intense green and yellow droplets, as anticipated due to increased charge screening. The observed transition from multiphase to single phase droplets between 150 and 300 mM KCl, as outer phase dissolves, is consistent with critical salt concentrations for R10/D10 and K10/D10 coacervates of approximately 1.5 M KCl and 250 mM KCl, respectively, in our previous study[8].

**Supplementary Discussion 3**
Lengths and hybridization status of RNAs are directly relevant to multivalency of molecules affecting partitioning and hybridization level of RNA in coacervate droplets, but other factors should be considered such as hydrogen bonding and cation-pi interactions as parts of RNA-peptide interactions. For example, ssRNA 20mer has the same level of multivalency as that of fully hybridized dsRNA 10mer. However, single-phase droplets had similar or slightly higher partitioning for ssRNA 20mer than that of dsRNA 10mer, indicating more intermolecular interactions rather than electrostatic interaction by multivalency of RNAs. Remarkably, pre-hybridized dsRNA 10mer shows significant greater change like 9-fold less partitioning in inner droplets of R10/K10/D10 and 3-fold increase in partitioning in outer droplets than that of ssRNA 20mer. This suggests that dsRNA has different levels of hybridization upon accumulation in each phase of multiphase droplets.

**Supplementary Discussion 4**
If any melting occurred in coacervates, [dsRNA]* in Fig. 2 could arise from a combination of duplexed and free forms of the monitored fluorescent strand. Thus, we further analyze to distinguish Cy3-RNA signal from actual dsRNA versus its non-hybridized form using estimated fraction of hybridization of RNA duplex from FRET values in Fig. 3. The estimated concentration of dsRNA ([dsRNA] in Fig. 4) and ssRNA ([ssRNA] in Fig. 4) from [dsRNA]* is plotted in Fig. 4 and in Supplementary Table 12. Here we compare the trend of Cy3-RNA levels of ssRNA in Fig. 2C, to estimated ssRNA concentrations in Fig. 4.

The 10-mer and 20-mer [ssRNA] in Fig. 2C was 10x and 30x higher in the inner phase than outer phase. In Fig. 4, the estimated [ssRNA] was 10-fold higher for the 10-mer in the inner phase than the outer phase of R10/K10/D10 (same as [ssRNA] trend in Fig. 2C) and 1.5-fold more concentrated for the 20-mer in the inner R10/K10/D10 coacervate droplets (less pronounced ssRNA partitioning in inner phase than [ssRNA] in Fig. 2C). While these values are statistically similar $\Delta G_{P,ss}$ for the ssRNA 10-mer, they are different for the ssRNA 20-mer in Supplementary Fig. 7B. We hypothesize that the dissociated, complementary ssRNA strands from initially-dsRNA partition somewhat less favorably into the inner coacervate phase of R10/K10/D10 due to the additional interactions with their complementary strands and perhaps also to the equilibria between free and duplexed forms across the coacervate phases.

**References**.

1 Schindelin, J. *et al.* Fiji: an open-source platform for biological-image analysis. *Nat. Methods* **9**, 676-682 (2012).
2 Fu, J., Fares, H. M. & Schlenoff, J. B. Ion-Pairing Strength in Polyelectrolyte Complexes. *Macromolecules* **50**, 1066-1074, doi:10.1021/acs.macromol.6b02445 (2017).
3 Pir Cakmak, F., Grigas, A. T. & Keating, C. D. Lipid vesicle-coated complex coacervates. *Langmuir* **35**, 7830-7840 (2019).

4 Perry, S. L., Li, Y., Priftis, D., Leon, L. & Tirrell, M. The effect of salt on the complex coacervation of vinyl polyelectrolytes. *Polymers* **6**, 1756-1772 (2014).
5 Pace, C. N., Grimsley, G. R. & Scholtz, J. M. Protein ionizable groups: pK values and their contribution to protein stability and solubility. *J. Biol. Chem.* **284**, 13285-13289 (2009).
6 Van der Gucht, J., Spruijt, E., Lemmers, M. & Stuart, M. A. C. Polyelectrolyte complexes: Bulk phases and colloidal systems. *J. Colloid Interface Sci.* **361**, 407-422 (2011).
7 Fisher, R. S. & Elbaum-Garfinkle, S. Tunable multiphase dynamics of arginine and lysine liquid condensates. *Nat. Commun.* **11**, 1-10 (2020).
8 Cakmak, F. P., Choi, S., Meyer, M. O., Bevilacqua, P. C. & Keating, C. D. Prebiotically-relevant low polyion multivalency can improve functionality of membraneless compartments. *Nat. Commun.* **11**, 5949, doi:10.1038/s41467-020-19775-w (2020).

**Supplementary Figures**

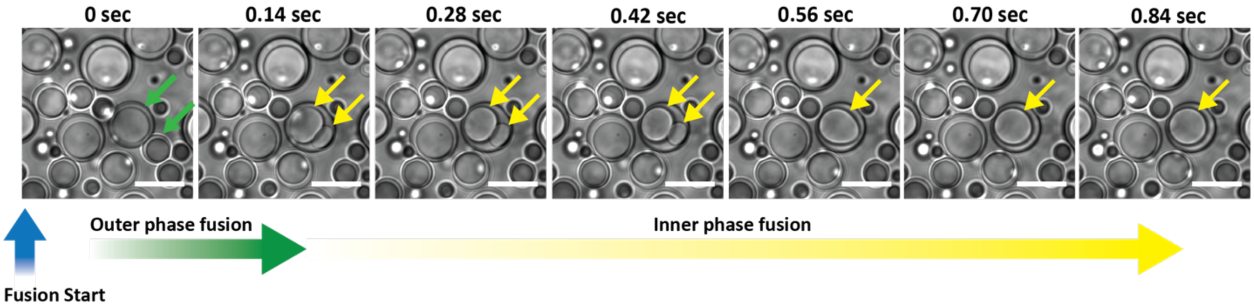


**Supplementary Figure 1.** Snapshots of fusion of R10/K10/D10 multiphase coacervate droplets. Two droplets indicated by the arrows in frame 1 (0 sec) become 1 droplet indicated by the arrow in frame 2 (0.14 sec), but for inner phase of R10/K10/D10 took several more frames to be fully fused into spherical shapes (from 0.14 sec to 0.84 sec). Thus, fusion of outer phase of R10/K10/D10 is faster than the fusion of inner phase of R10/K10/D10. Scale bars = 20 $\mu M$.

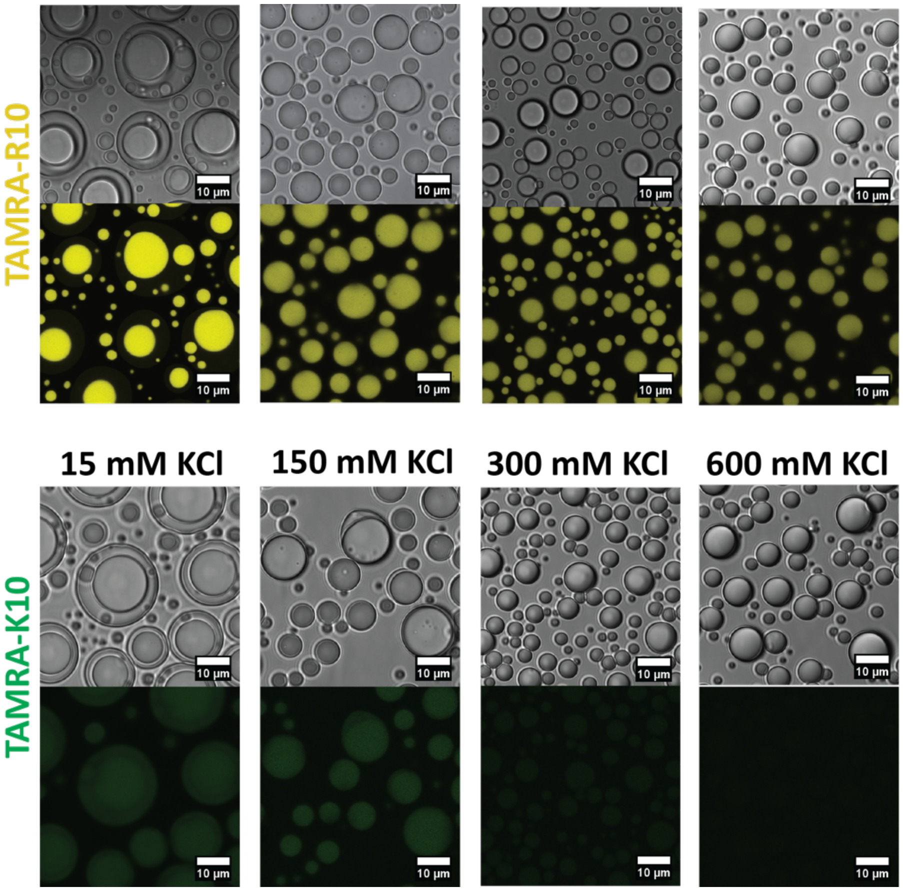


**Supplementary Figure 2.** Images of multiphase complex coacervates of R10/K10/D10 as salt concentration is increased. Confocal microscope settings are equal between images of different salt concentrations.

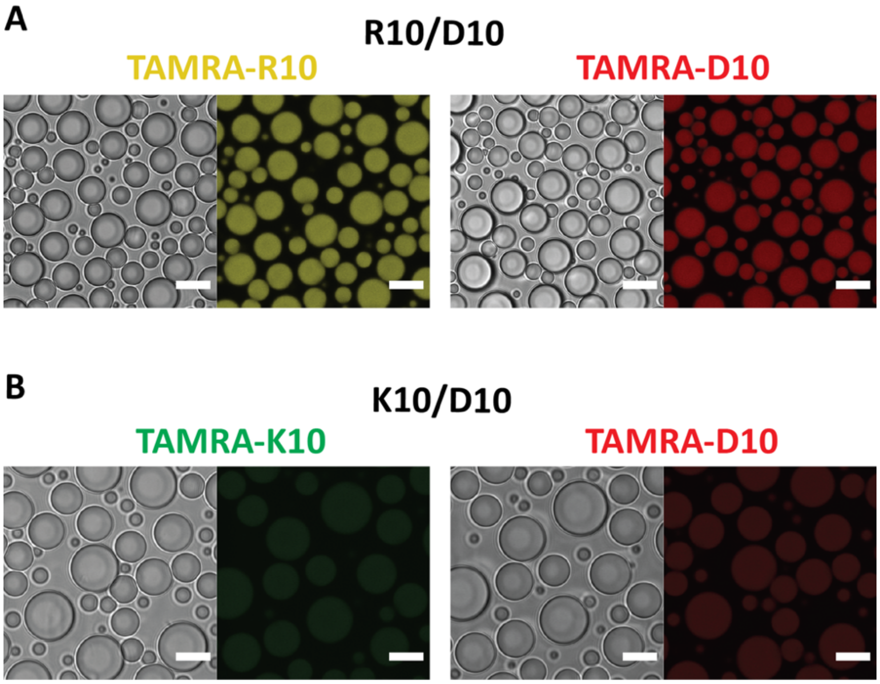


**Supplementary Figure 3.** Images of oligopeptides partitioning in single-phase coacervate droplets of (A) R10/D10 and (B) K10/D10. Scale bars = 10 $\mu m$.

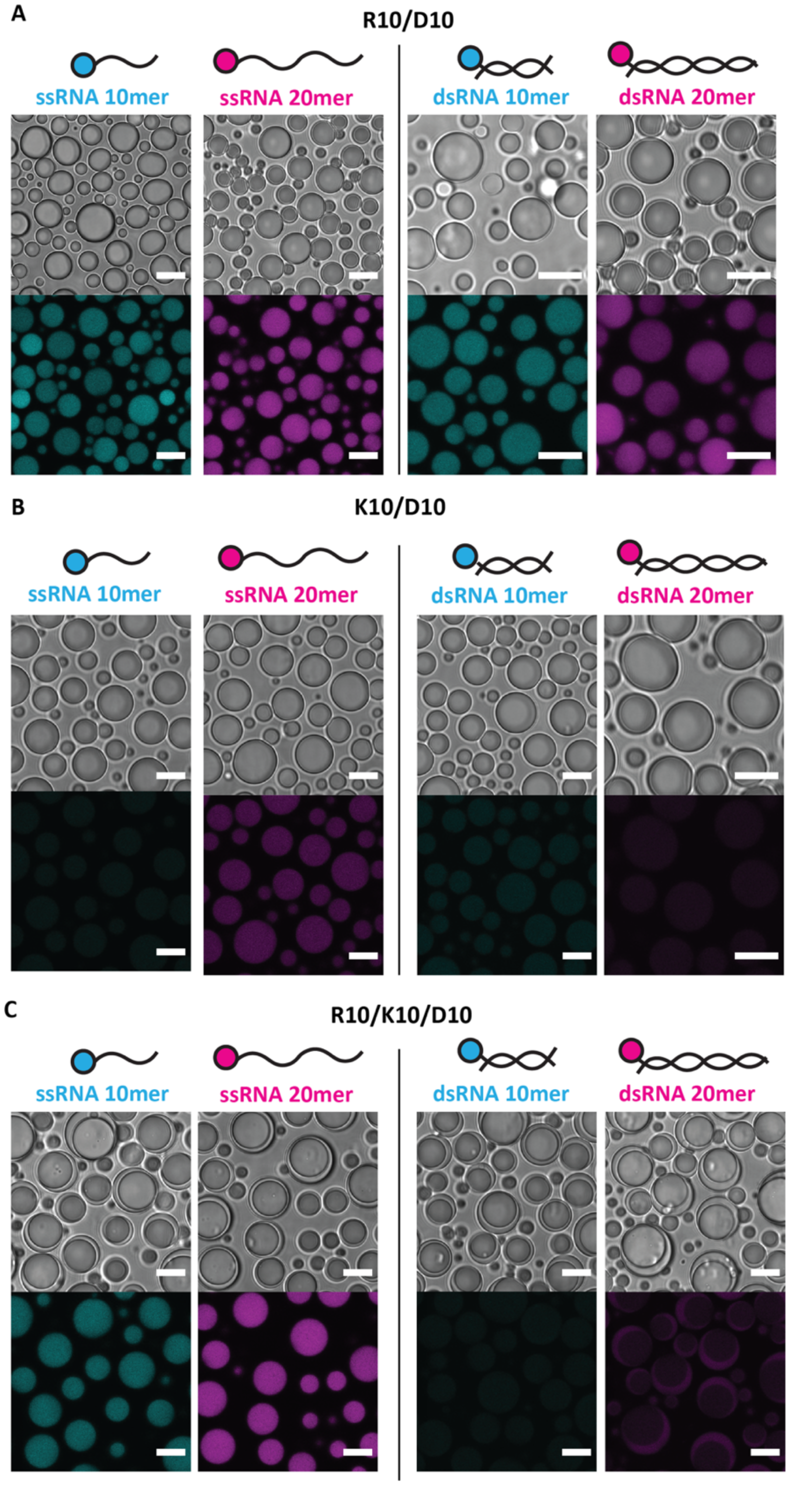


**Supplementary Figure 4.** Images of RNA partitioning in (A) R10/D10 coacervates, (B) K10/D10 coacervates and (C) R10/K10/D10 coacervates. Confocal settings are equal among images. Scale bars = 10 μm. These data were used to construct calibration plots in Fig. 2A and C.

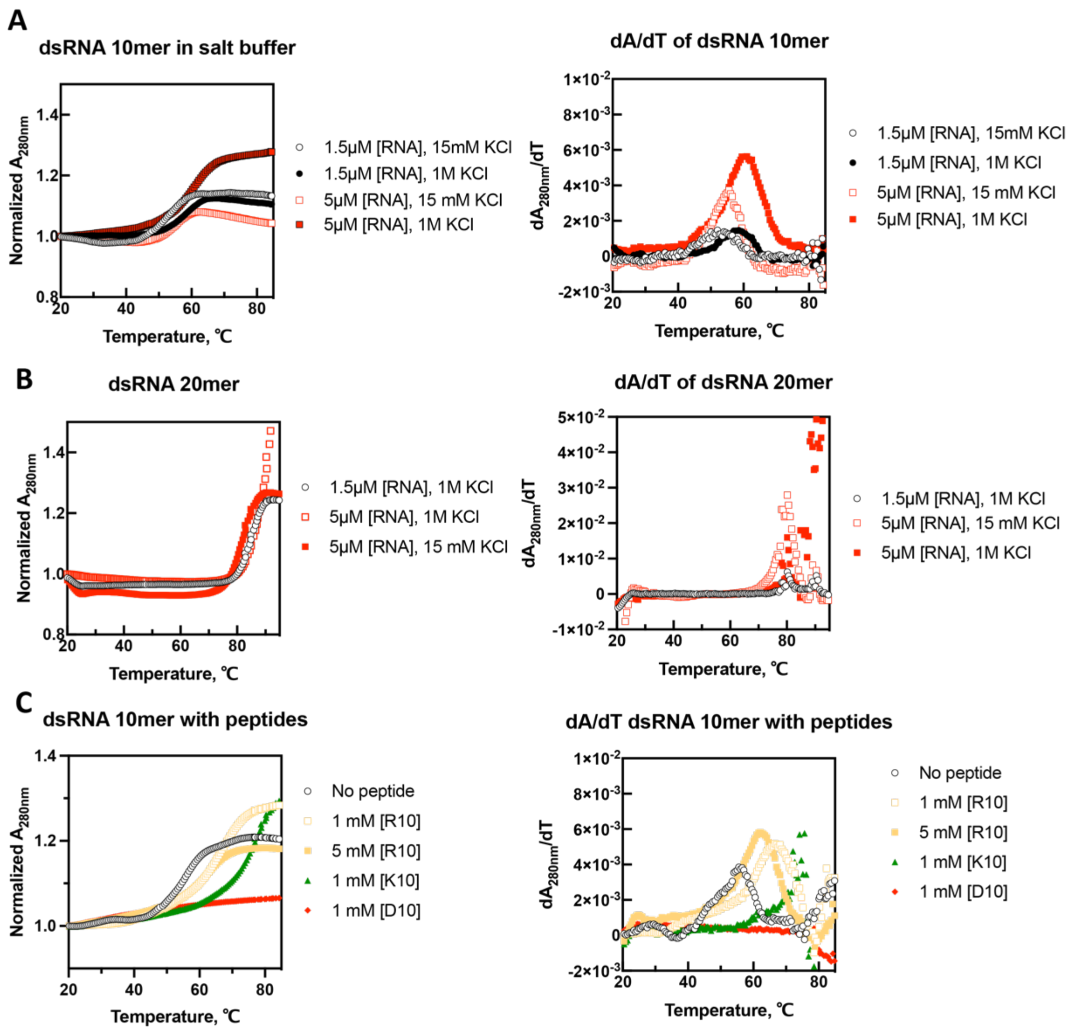


**Supplementary Figure 5.** UV-Vis melting curves and the first derivative of curves of dsRNA 10mer and 20mer in various solution conditions. Melting of varying lengths of RNA duplex and salt concentrations of (A) dsRNA 10mer and (B) dsRNA 20mer. All conditions in (A) and (B) contain 0.5 mM $MgCl_2$ in addition to KCl stated in plots, and are at pH 8.2 ± 0.1. (C) Melting of RNA 10mer duplexes in presence of peptides in 15 mM KCl, 0.5 mM $MgCl_2$ and pH 8.2 ± 0.1. Melting of dsRNA 20mer with peptides could not be obtained since the melting temperatures are beyond the range of temperature we can measure. UV-Vis melting of prehybridized dsRNA is analyzed by a diode array OLIS spectrometer starting from 20 °C and heated to 90 ~ 95 °C with 0.5 °C increment and 30 seconds of equilibration in between. Melting curves are 2$^{nd}$ order smoothing done for 11 neighbors using GraphPad Prism 9.

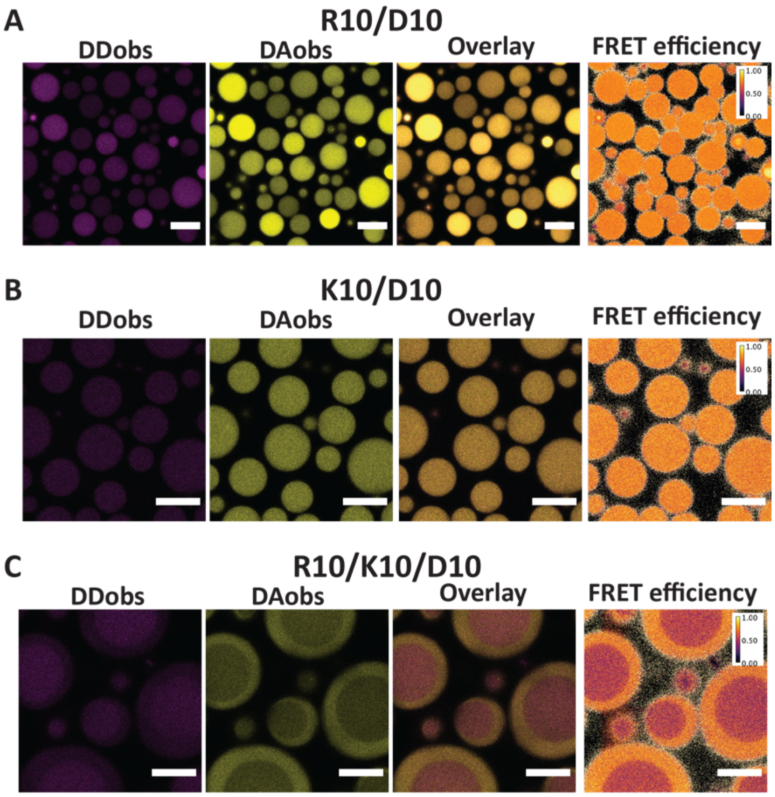


**Supplementary Figure 6.** Fluorescence images and FRET efficiency plots of dsRNA 20mer in (A) R10/D10 and (B) K10/D10 single-phase coacervates, and R10/K10/D10 multiphase coacervates. Scale bars = 10 µm.

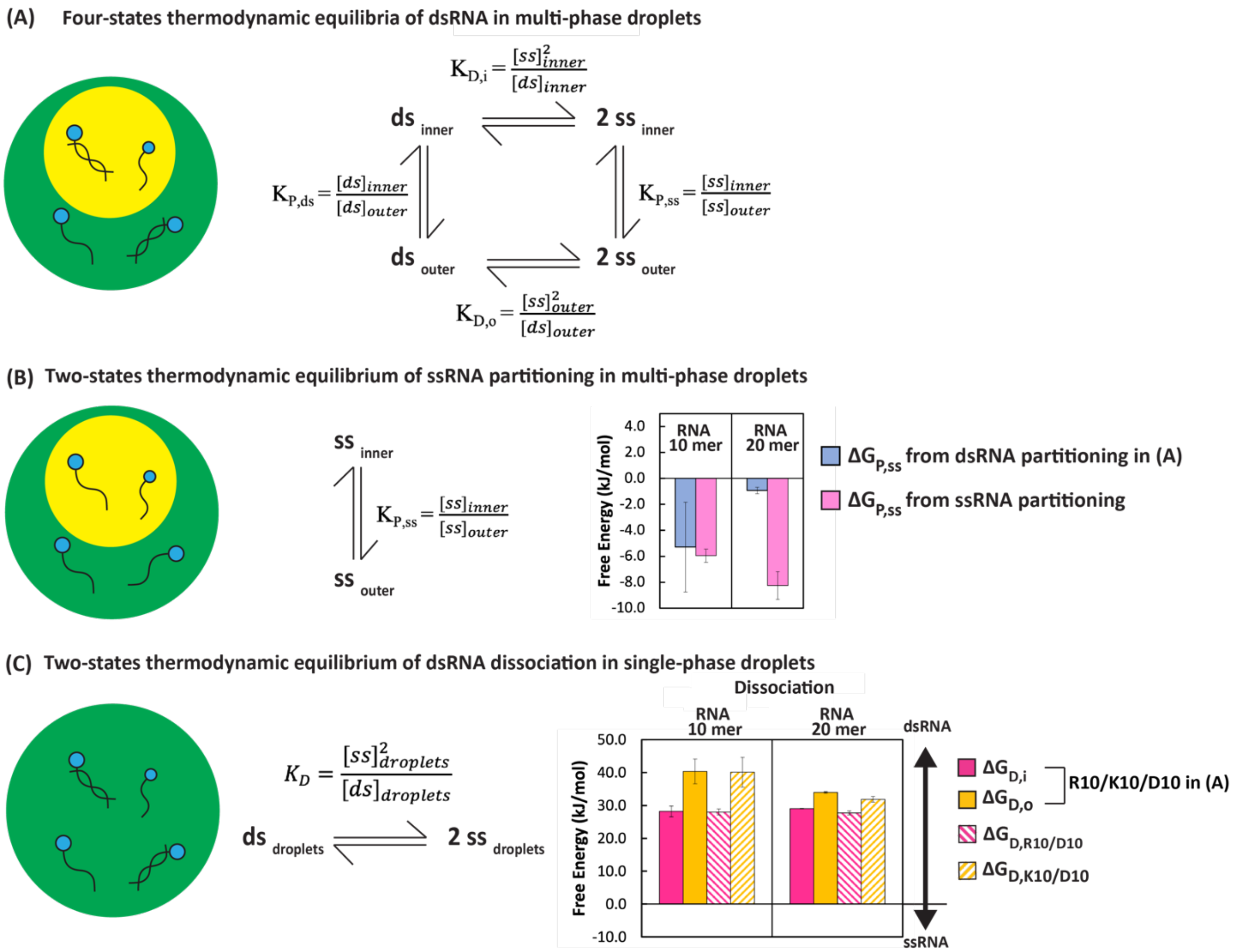


**Supplementary Figure 7.** (A) Thermodynamic equilibria of four states of RNA in inner and outer phase of R10/K10/D10. (B) Thermodynamic equilibrium of two states of ssRNA partitioning in multiphase droplets. Bar graphs are plots of Gibbs free energy changes of ssRNA partitioning from ssRNA partitioning experiments (Fig. 2 A, C) to compare to that from dsRNA partitioning experiments (as in (A), same values as in Fig. 5B). (C) Thermodynamic equilibrium of two states of RNA in single-phase of droplets. Bar graphs are plots of Gibbs free energy change in dissociation of dsRNA into ssRNA in inner and outer droplets of R10/K10/D10 (as in (A), same values as in Fig. 5D) compared to that of R10/D10 single-phase droplets or K10/D10 single-phase droplets. Values are in Supplementary Table 13, 14 and 15.

### Supplementary Tables

**Supplementary Table 1.** Final total concentration of fluorescently unlabeled and labeled-peptides in samples for each measurement.

| | **[R10]** | | **[K10]** | | **[D10]** | | **Dilution factor of labeled peptide to non-labeled peptide** |
|---|---|---|---|---|---|---|---|
| | Peptide | Labeled Peptide | Peptide | Labeled Peptide | Peptide | Labeled Peptide | |
| **R10/D10** | | | | | | | |
| RNA characterization | 2.000 mM | N/A | N/A | | 2.000 mM | N/A | N/A |
| [R10] quantification | 1.998 mM | 0.002 mM | N/A | | 2.000 mM | N/A | 1000 |
| [D10] quantification | 2 mM | N/A | N/A | | 1.998 mM | 0.002 mM | 1000 |
| **K10/D10** | | | | | | | |
| RNA characterization | N/A | | 2.000 mM | N/A | 2.000 mM | N/A | N/A |
| [K10] quantification | N/A | | 1.998 mM | 0.002 mM | 2 mM | N/A | 1000 |
| [D10] quantification | N/A | | 2 mM | N/A | 1.998 mM | 0.002 mM | 1000 |
| **R10/K10/D10** | | | | | | | |
| RNA characterization | 2.000 mM | N/A | 2.000 mM | N/A | 4.000 mM | N/A | N/A |
| [R10] quantification | 1.998 mM | 0.002 mM | 2.000 mM | N/A | 4.000 mM | N/A | 1000 |
| [K10] quantification | 2.000 mM | N/A | 1.998 mM | 0.002 mM | 4.000 mM | N/A | 1000 |
| [D10] quantification | 2.000 mM | N/A | 2.000 mM | N/A | 3.998 mM | 0.002 mM | 2000 |

**Supplementary Table 2.** Concentrations of R10, K10 and D10 in multiphase complex coacervate droplets. These are plotted in Figure 1 (D). Errors bars are calculated from standard deviation of intensity values by the propagation of errors with calibration curves.

| | **$[R10]_{monomer}$ (mM)** | **$[K10]_{monomer}$ (mM)** | **$[D10]_{monomer}$ (mM)** | **$(+)_{total, monomer}/(-)_{monomer}$** | **$[R]_{monomer}/[K]_{monomer}$** |
|---|---|---|---|---|---|
| **R10/D10 droplets** | 710 ± 96 | N/A | 731 ± 43 | 0.97 ± 0.07 | 6.5 ± 0.6 |
| **K10/D10 droplets** | N/A | 110 ± 34 | 371 ± 16 | 0.30 ± 0.03 | |
| **R10/K10/D10 inner droplets** | 755 ± 28 | 68 ± 5 | 911 ± 38 | 0.90 ± 0.05 | 11.1 ± 1.0 |
| **R10/K10/D10 outer droplets** | 26 ± 2 | 28 ± 4 | 21 ± 9 | 2.59 ± 0.43 | 0.9 ± 0.2 |

**Supplementary Table 3.** P values of monomer concentrations in coacervate phases done by two-sided t-test with unequal variance.

| Sample 1 | Sample 2 | p-values |
|---|---|---|
| **[R10]** | | |
| R10/D10 | R10/K10/D10 inner | P = 0.0421, * |
| R10/D10 | R10/K10/D10 outer | P < 0.0001, **** |
| R10/K10/D10 inner | R10/K10/D10 outer | P < 0.0001, **** |
| **[K10]** | | |
| K10/D10 | R10/K10/D10 inner | P < 0.0001, **** |
| K10/D10 | R10/K10/D10 outer | P < 0.0001, **** |
| R10/K10/D10 inner | R10/K10/D10 outer | P < 0.0001, **** |
| **[D10]** | | |
| R10/D10 | K10/D10 | |
| R10/D10 | R10/K10/D10 inner | P < 0.0001, **** |
| R10/D10 | R10/K10/D10 outer | P = 0.0751, ns |
| K10/D10 | R10/K10/D10 inner | P < 0.0001, **** |
| K10/D10 | R10/K10/D10 outer | P < 0.0001, **** |
| R10/K10/D10 inner | R10/K10/D10 outer | P < 0.0001, **** |

**Supplementary Table 4.** Sequences of RNA pairs used in this paper. [a]

| Label | Sequence 1 (5' to 3') | Sequence 2 (5' to 3') |
|---|---|---|
| **RNA 10mer** | | |
| FRET sample or dsRNA control | ACCUUGUUCC[Cy3] | [Cy5]GGAACAAGGU |
| Donor sample | ACCUUGUUCC[Cy3] | GGAACAAGGU |
| Acceptor sample | ACCUUGUUCC | [Cy5]GGAACAAGGU |
| ssRNA control | [Cy3]GGAACAAGGU | [Cy5]GGAACAAGGU |
| **RNA 20mer** | | |
| FRET sample or dsRNA control | AUCUCGCUCUACCUUGUUCC[Cy3] | [Cy5]GGAACAAGGUAGAGCGAGAU |
| Donor sample | AUCUCGCUCUACCUUGUUCC[Cy3] | GGAACAAGGUAGAGCGAGAU |
| Acceptor sample | AUCUCGCUCUACCUUGUUCC | [Cy5]GGAACAAGGUAGAGCGAGAU |
| ssRNA control | [Cy3]GGAACAAGGUAGAGCGAGAU | [Cy5]GGAACAAGGUAGAGCGAGAU |

[a] The first three rows of each section are a pair of complementary sequences. Note that the 20mers start with the same 10 nucleotides as the 10mers. Self-complementary is avoided by having one strand (Sequence 1) be pyrimidine-rich and the other strand (Sequence 2) be purine-rich. All duplexes have 50% GC content, which is spread out along the length of each RNA. The Cy3 and Cy5 labels end up on the same end of the duplex.

**Supplementary Table 5.** Concentration of labeled RNA in multiphase complex coacervate droplets. Unit of concentration is μM.

| | **Initial forms of RNAs mixed with coacervate samples** | | | |
|---|---|---|---|---|
| | **ssRNA 10mer** | **ssRNA 20mer** | **dsRNA 10mer** | **dsRNA 20mer** |
| **R10/D10 droplets** | 44.8 ± 12.1 | 68.1 ± 11.4 | 42.8 ± 2.6 | 53.8 ± 15.1 |
| **K10/D10 droplets** | 10.6 ± 0.9 | 26.3 ± 5.5 | 10.2 ± 1.2 | 10.9 ± 7.3 |
| **R10/K10/D10 inner droplets** | 43.4 ± 2.9 | 66.6 ± 7.5 | 7.1 ± 2.7 | 8.6 ± 0.8 |
| **R10/K10/D10 outer droplets** | 4.0 ± 0.9 | 2.3 ± 0.9 | 7.6 ± 3.9 | 20.6 ± 2.1 |

The term of "Initial forms of RNAs" indicates that dsRNAs might be dissociated when they are in coacervate phases.

**Supplementary Table 6.** P-values to compare RNAs of different lengths. Two-sided t-test with unequal variance was performed between Sample 1 and Sample 2.

| Sample 1 | Sample 2 | p-values |
|---|---|---|
| **ssRNA 10mer** | **ssRNA 20mer** | |
| R10/D10 | R10/D10 | P<0.0001, **** |
| K10/D10 | K10/D10 | P<0.0001, **** |
| R10/K10/D10 in | R10/K10/D10 in | P<0.0001, **** |
| R10/K10/D10 out | R10/K10/D10 out | P<0.0001, **** |
| **dsRNA 10mer** | **dsRNA 20mer** | |
| R10/D10 | R10/D10 | P<0.0001, **** |
| K10/D10 | K10/D10 | P<0.0001, **** |
| R10/K10/D10 in | R10/K10/D10 in | P<0.0001, **** |
| R10/K10/D10 out | R10/K10/D10 out | P=0.0007, *** |

**Supplementary Table 7.** P-values to compare ssRNA and dsRNA of same length of RNA. Two-sided t-test with unequal variance was performed between Sample 1 and Sample 2.

| Sample 1 | Sample 2 | p-values |
|---|---|---|
| **ssRNA 10mer** | **dsRNA 10mer** | |
| R10/D10 | R10/D10 | P = 0.2755, ns |
| K10/D10 | K10/D10 | P = 0.0944, ns |
| R10/K10/D10 inner | R10/K10/D10 inner | P<0.0001, **** |
| R10/K10/D10 outer | R10/K10/D10 outer | P<0.0001, **** |
| **ssRNA 20mer** | **dsRNA 20mer** | |
| R10/D10 | R10/D10 | P<0.0001, **** |
| K10/D10 | K10/D10 | P<0.0001, **** |
| R10/K10/D10 inner | R10/K10/D10 inner | P<0.0001, **** |
| R10/K10/D10 outer | R10/K10/D10 outer | P<0.0001, **** |

**Supplementary Table 8.** P-values to compare same RNA in different phases. Two-sided t-test with unequal variance was performed between Sample 1 and Sample 2.

| Sample 1 | Sample 2 | p-values |
|---|---|---|
| **ssRNA 10mer** | | |
| R10/D10 | K10/D10 | P < 0.0001, **** |
| R10/D10 | R10/K10/D10 inner | P = 0.4601, ns |
| R10/D10 | R10/K10/D10 outer | P < 0.0001, **** |
| **K10/D10** | R10/K10/D10 inner | P < 0.0001, **** |
| K10/D10 | R10/K10/D10 outer | P < 0.0001, **** |
| R10/K10/D10 inner | R10/K10/D10 outer | P < 0.0001, **** |
| **ssRNA 20mer** | | |
| R10/D10 | K10/D10 | P < 0.0001, **** |
| R10/D10 | R10/K10/D10 inner | P = 0.4513, ns |
| R10/D10 | R10/K10/D10 outer | P < 0.0001, **** |
| K10/D10 | R10/K10/D10 inner | P < 0.0001, **** |
| K10/D10 | R10/K10/D10 outer | P < 0.0001, **** |
| R10/K10/D10 inner | R10/K10/D10 outer | P < 0.0001, **** |
| **dsRNA 10mer** | | |
| R10/D10 | K10/D10 | P < 0.0001, **** |
| R10/D10 | R10/K10/D10 inner | P < 0.0001, **** |
| R10/D10 | R10/K10/D10 outer | P < 0.0001, **** |
| K10/D10 | R10/K10/D10 inner | P < 0.0001, **** |
| K10/D10 | R10/K10/D10 outer | P < 0.0001, **** |
| R10/K10/D10 inner | R10/K10/D10 outer | P = 0.5087, ns |
| **dsRNA 20mer** | | |
| R10/D10 | K10/D10 | P < 0.0001, **** |
| R10/D10 | R10/K10/D10 inner | P < 0.0001, **** |
| R10/D10 | R10/K10/D10 outer | P < 0.0001, **** |
| K10/D10 | R10/K10/D10 inner | P = 0.0430, * |
| K10/D10 | R10/K10/D10 outer | P < 0.0001, **** |
| R10/K10/D10 inner | R10/K10/D10 outer | P<0.0001, **** |

**Supplementary Table 9.** Correction values for FRET efficiency calculations.

| | $\alpha$ | $\beta$ |
|---|---|---|
| **RNA 10mer** | | |
| R10/K10/D10 inner droplets | 0.0681 ± 0.0165 | 0.0775 ± 0.0132 |
| R10/K10/D10 outer droplets | 0.0655 ± 0.0169 | 0.0744 ± 0.0124 |
| R10/D10 droplets | 0.0532 ± 0.0055 | 0.0770 ± 0.0033 |
| K10/D10 droplets | 0.0770 ± 0.0033 | 0.0944 ± 0.0015 |
| dsRNA buffer control | 0.0585 ± 0.0176 | 0.0659 ± 0.0003 |
| ssRNA buffer control | 0.0686 ± 0.0150 | 0.0504 ± 0.0033 |
| **RNA 20mer** | | |
| R10/K10/D10 inner droplets | 0.0609 ± 0.0066 | 0.0525 ± 0.0070 |
| R10/K10/D10 outer droplets | 0.0607 ± 0.0101 | 0.0560 ± 0.0022 |
| R10/D10 droplets | 0.0603 ± 0.0069 | 0.0575 ± 0.0121 |
| K10/D10 droplets | 0.0625 ± 0.0066 | 0.0574 ± 0.0008 |
| dsRNA buffer control | 0.0141 ± 0.0095 | 0.0741 ± 0.0031 |
| ssRNA buffer control | 0.0686 ± 0.0150 | 0.0504 ± 0.0033 |

**Supplementary Table 10.** FRET efficiency values. Errors bars are calculated from standard deviation of three measurements of three samples.

| | **FRET efficiency** | **Normalized FRET efficiency** |
|---|---|---|
| **RNA 10mer** | | |
| R10/K10/D10 inner droplets | 0.38 ± 0.04 | 0.54 |
| R10/K10/D10 outer droplets | 0.68 ± 0.06 | 0.95 |
| R10/D10 droplets | 0.55 ± 0.08 | 0.77 |
| K10/D10 droplets | 0.68 ± 0.08 | 0.95 |
| dsRNA buffer | 0.72 ± 0.08 | 1.00 |
| ssRNA buffer | 0.00 ± 0.00 | 0.00 |
| **RNA 20mer** | | |
| R10/K10/D10 inner droplets | 0.52 ± 0.01 | 0.62 |
| R10/K10/D10 outer droplets | 0.75 ± 0.01 | 0.89 |
| R10/D10 droplets | 0.66 ± 0.05 | 0.79 |
| K10/D10 droplets | 0.66 ± 0.06 | 0.78 |
| dsRNA buffer | 0.84 ± 0.10 | 1.00 |
| ssRNA buffer | 0.00 ± 0.00 | 0 |

**Supplementary Table 11.** P-values to compare FRET efficiency of coacervate phases to FRET efficiency of dsRNA in salt buffer. Two-sided t-test with unequal variance was performed between Sample 1 and Sample 2.

| **Sample 1** | **Sample 2** | **p-values** |
|---|---|---|
| **RNA 10mer** | | |
| dsRNA 10mer in buffer | ssRNA 10mer in buffer | P<0.0001, **** |
| dsRNA 10mer in buffer | R10/D10 | P= 0.0003 , *** |
| dsRNA 10mer in buffer | K10/D10 | P=0.3881, ns |
| dsRNA 10mer in buffer | R10/K10/D10 inner | P<0.0001, **** |
| dsRNA 10mer in buffer | R10/K10/D10 outer | P=0.2550, ns |
| R10/D10 | K10/D10 | P= 0.0029, ** |
| R10/D10 | R10/K10/D10 inner | P=0.0001, *** |
| R10/D10 | R10/K10/D10 outer | P=0.0014, ** |
| K10/D10 | R10/K10/D10 inner | P<0.0001, **** |
| K10/D10 | R10/K10/D10 outer | P=0.9259, ns |
| **R10/K10/D10 inner** | R10/K10/D10 outer | P<0.0001, **** |
| **RNA 20mer** | | |
| dsRNA 20mer in buffer | ssRNA 10mer in buffer | P<0.0001, **** |
| dsRNA 20mer in buffer | R10/D10 | P= 0.0004, *** |
| dsRNA 20mer in buffer | K10/D10 | P= 0.0004 , *** |
| dsRNA 20mer in buffer | R10/K10/D10 in | P<0.0001, **** |
| dsRNA 20mer in buffer | R10/K10/D10 outer | P= 0.0219, * |
| R10/D10 | K10/D10 | P= 0.8684, ns |
| R10/D10 | R10/K10/D10 inner | P<0.0001, **** |
| R10/D10 | R10/K10/D10 outer | P=0.0011, ** |
| K10/D10 | R10/K10/D10 inner | P<0.0001, **** |
| K10/D10 | R10/K10/D10 outer | P= 0.0020, ** |
| R10/K10/D10 inner | R10/K10/D10 outer | P<0.0001, **** |

**Supplementary Table 12.** Estimated dsRNA and ssRNA concentrations from pre-hybridized dsRNA partitioning experiment using normalized FRET efficiency in coacervate droplets.

| | **[dsRNA], $\mu M$** | **[ssRNA], $\mu M$** |
|---|---|---|
| **RNA 10mer** | | |
| R10/K10/D10 inner droplets | 3.80 ± 1.45 | 6.58 ± 2.52 |
| R10/K10/D10 outer droplets | 7.17 ± 3.72 | 0.78 ± 1.05 |
| R10/D10 droplets | 32.73 ± 3.83 | 20.10 ± 6.70 |
| K10/D10 droplets | 9.72 ± 1.44 | 0.95 ± 1.74 |
| **RNA 20mer** | | |
| R10/K10/D10 inner droplets | 5.31 ± 0.11 | 6.58 ± 0.18 |
| R10/K10/D10 outer droplets | 18.43 ± 0.42 | 4.54 ± 0.44 |
| R10/D10 droplets | 46.19 ± 3.68 | 25.22 ± 6.07 |
| K10/D10 droplets | 8.55 ± 1.23 | 4.71 ± 1.42 |

**Supplementary Table 13.** Thermodynamic parameters of partitioning ($K_{P,ds}$, $K_{P,ss}$, $\Delta G_{P,ds}$, and $\Delta G_{P,ss}$) and dissociation of dsRNA ($K_{D,i}$, $K_{D,o}$, $\Delta G_{D,i}$, and $\Delta G_{D,o}$) from pre-hybridized dsRNA partitioning in R10/K10/D10. Errors bars are calculated by error propagation methods using standard deviation of three measurements of 45 samples of partitioning experiment and 9 samples from FRET experiment of three independent trials.

| | RNA 10mer | RNA 20mer |
|---|---|---|
| **$K_{P,ds}$** | $(5.30 \pm 3.41) \times 10^{-1}$ | $(2.88 \pm 0.09) \times 10^{-1}$ |
| **$K_{P,ss}$** | $(8.44 \pm 1.18) \times 10^{+1}$ | $(1.45 \pm 0.14) \times 10^{0}$ |
| **$K_{D,i}$** | $(1.14 \pm 0.76) \times 10^{-5}$ | $(8.14 \pm 0.33) \times 10^{-6}$ |
| **$K_{D,o}$** | $(0.85 \pm 1.30) \times 10^{-7}$ | $(1.12 \pm 0.13) \times 10^{-6}$ |
| **$\Delta G_{P,ds}$, kJ/mol** | $(1.57 \pm 1.60) \times 10^{0}$ | $(3.08 \pm 0.08) \times 10^{0}$ |
| **$\Delta G_{P,ss}$, kJ/mol** | $(-5.28 \pm 3.46) \times 10^{0}$ | $(-9.20 \pm 0.25) \times 10^{-1}$ |
| **$\Delta G_{D,i}$, kJ/mol** | $(2.82 \pm 0.16) \times 10^{+1}$ | $(2.90 \pm 0.01) \times 10^{+1}$ |
| **$\Delta G_{D,o}$, kJ/mol** | $(4.03 \pm 0.38) \times 10^{+1}$ | $(3.40 \pm 0.03) \times 10^{+1}$ |

**Supplementary Table 14.** Thermodynamic parameters of dissociation of dsRNA in coacervate droplets ($K_{D,I}$, and $\Delta G_{D,I}$) from pre-hybridized dsRNA partitioning in R10/D10 or K10/D10. Errors bars are calculated by error propagation methods using standard deviation of 45 samples of partitioning experiments and 9 samples from FRET experiment of three independent trials.

| | RNA 10mer | RNA 20mer |
|---|---|---|
| **R10/D10 droplets** | | |
| **$K_{D,i}$** | $(1.23 \pm 0.11) \times 10^{-5}$ | $(1.38 \pm 0.37) \times 10^{-5}$ |
| **$\Delta G_{D,i}$, kJ/mol** | $(2.80 \pm 0.09) \times 10^{+1}$ | $(2.78 \pm 0.07) \times 10^{+1}$ |
| **K10/D10 droplets** | | |
| **$K_{D,i}$** | $(9.37 \pm 17.2) \times 10^{-8}$ | $(2.60 \pm 0.94) \times 10^{-6}$ |
| **$\Delta G_{D,i}$, kJ/mol** | $(4.01 \pm 0.45) \times 10^{+1}$ | $(3.19 \pm 0.09) \times 10^{+1}$ |

**Supplementary Table 15.** Thermodynamic parameters from ssRNA partitioning ([ssRNA] in Fig. 2) from outer phase to inner phase in R10/K10/D10 ($K_{P,ss}$ and $\Delta G_{P,ss}$). Errors bars are calculated by error propagation methods using standard deviation of 45 samples of partitioning experiments and 9 samples from FRET experiment of three independent trials.

| | RNA 10mer | RNA 20mer |
|---|---|---|
| **$K_{P,ss}$** | $(1.11 \pm 0.23) \times 10^{+1}$ | $(2.17 \pm 1.09) \times 10^{+1}$ |
| **$\Delta G_{P,ss}$, kJ/mol** | $(-5.95 \pm 0.51) \times 10^{0}$ | $(-8.25 \pm 1.07) \times 10^{0}$ |
